# Potential for transmission of *Elizabethkingia anophelis* by *Aedes albopictus* and the role of microbial interactions in Zika virus competence

**DOI:** 10.1101/702464

**Authors:** MG Onyango, AF Payne, J Stout, C Dieme, L Kuo, LD Kramer, AT Ciota

## Abstract

*Elizabethkingia anophelis* has been the cause of four outbreaks with significant morbidity and mortality. Its transmission routes remain unknown and no point source of infection has been identified. Here we show that *E. anophelis* can be found in the saliva of *Aedes* mosquitoes, suggesting the novel possibility of vector-borne transmission of this bacterium. We additionally characterized diverse microbial communities in *Aedes* midguts, salivary glands and saliva. To the best of our knowledge, this represents the first description of the microbiome of *Aedes* saliva. Further, we demonstrate that increased abundance of *E. anophelis* is associated with decreased susceptibility and replication of Zika virus (ZIKV) in the midgut of *Aedes* mosquitoes, suggesting a novel transmission barrier for arboviruses transmitted by *Aedes* mosquitoes. Together, these results demonstrate the complex relationships between the mosquito, the midgut microbial community and arboviruses and offer insights into the epidemiology and control of emerging bacterial and viral pathogens.

**Author Summary:** *Elizabethkingia anophelis* has in the recent past caused outbreaks different parts of the world resulting both in morbidity and mortality. Until now, to the best of our knowledge, no study has been able to demonstrate that this bacterium can be transmitted by mosquitoes. We have demonstrated for the first time that *Elizabethkingia anophelis* is present in the saliva of both infected and non-infected *Aedes* mosquitoes. Further, we have shown that it confers an inhibitory effect on Zika virus establishment in the midguts of *Aedes* mosquitoes. Together, these results potentially display the potential for vector borne transmission of *E. anophelis* as well as a novel transmission barrier of ZIKV. Lastly, we have for the first time characterized salivary microbes of *Aedes* mosquitoes necessitating the investigation of the impact of salivary microbes in severity of disease in vertebrate hosts.

## Introduction

*Elizabethkingia* is a gram-negative rod-like, aerobic, non-fermenting, non-motile and non-spore forming bacteria widely distributed in natural environments, including soil and fresh water as well as hospital environments [1]. It was originally called Flavobacterium or belonged to CDC group IIa but later reclassified as Chryseobacterium in 1994 [2]. On the basis of shared phenotypic and phylogenetic characteristics, *Chryseobacterium meningosepticum* and *C. miricola* were re-classified to a new genus, *Elizabethkingia* [3]. The most clinically important species of the genus *Elizabethkingia* include *E. anophelis, E. meningoseptica and E. miricola* [4]. *E. anophelis*, originally isolated from the midgut of *Anopheles gambiae* [5] has been the cause of moderate to large-scale outbreaks resulting in morbidity and mortality [6–10] as well as clinical cases across different geographical settings [1,4,11,12]. *E. anophelis* is associated with clinically significant infections and high mortality [4] relative to *E. meningoseptica* [9]. The genus *Elizabethkingia* is prevalent in *Aedes* [13–15] and *Anopheles* [16–18] species of mosquitoes and has been detected in both field-caught and laboratory-reared mosquitoes in Africa, Europe and North America [15,18,19]. The microbe can be transmitted among mosquitoes vertically, transstadially and horizontally [13,17]. In humans, evidence for perinatal vertical transmission of *E. anophelis* has been identified and associated with neonatal meningitis [11]. While some infections are thought to be nosocomial and mechanical ventilation has been identified as a risk factor for transmission in hospital settings [1], transmission routes have not been identified in a number of recent individual cases or clusters. The transition of *Elizabethkingia* from a common environmental bacterial species to a human pathogen of clinical significance, as well as the previous identification of *Elizabethkingia* in a number of mosquito species, prompted us to assess the abundance and tropism of *Elizabethkingia* bacteria in *Aedes* mosquitoes. We identified *Elizabethkingia* in midguts, salivary glands and saliva of *Ae. albopictus*. Sequencing of *Elizabethkingia* identified in the salivary gland demonstrated 100% homology to *E. anophelis*.

In order to assess the relative abundance of *Elizabethkingia* within microbial communities in these tissues, we additionally performed 16S *rRNA* sequencing. Further, to better define interactions with arboviruses vectored by *Aedes* mosquitoes, we assessed the correlation between *Elizabethkingia* and other microbes with Zika virus (ZIKV) in the midgut of *Ae. albopictus*. We measured a negative correlation between *E. anophelis* and ZIKV in the midgut and in mosquito cell culture. These data together demonstrate the complex interactions between microbial communities and arboviruses that could have important implications for the emergence and transmission of both viral and bacterial pathogens.

## Results

### Microbial profiles in *Aedes albopictus* are tissue-specific

Illumina MiSeq 16S *rRNA* sequencing resulted in 2 369 009 reads. The number of reads for individual samples ranged from 11 925 - 105 039 with a mean of 67 334. Both the positive spike-in (PhiX) and negative controls (water and non-template control) had limited number of sequences reads, ranging from 1620 – 2823, and hence were discarded in downstream analysis. In order to control for alpha and beta diversity biases, each sample was rarefied to a total frequency of 30 000 reads, hence two samples were discarded from downstream analysis. A total of 154 OTUs were identified in this study, 59% were Bacteroidetes, 40% Proteobacteria and 1% Cyanobacteria. In order to visualize the distribution of bacterial genera across individual tissues, an additional cut-off read frequency of 7 000 was applied (Figure 1).

**Figure 1.**
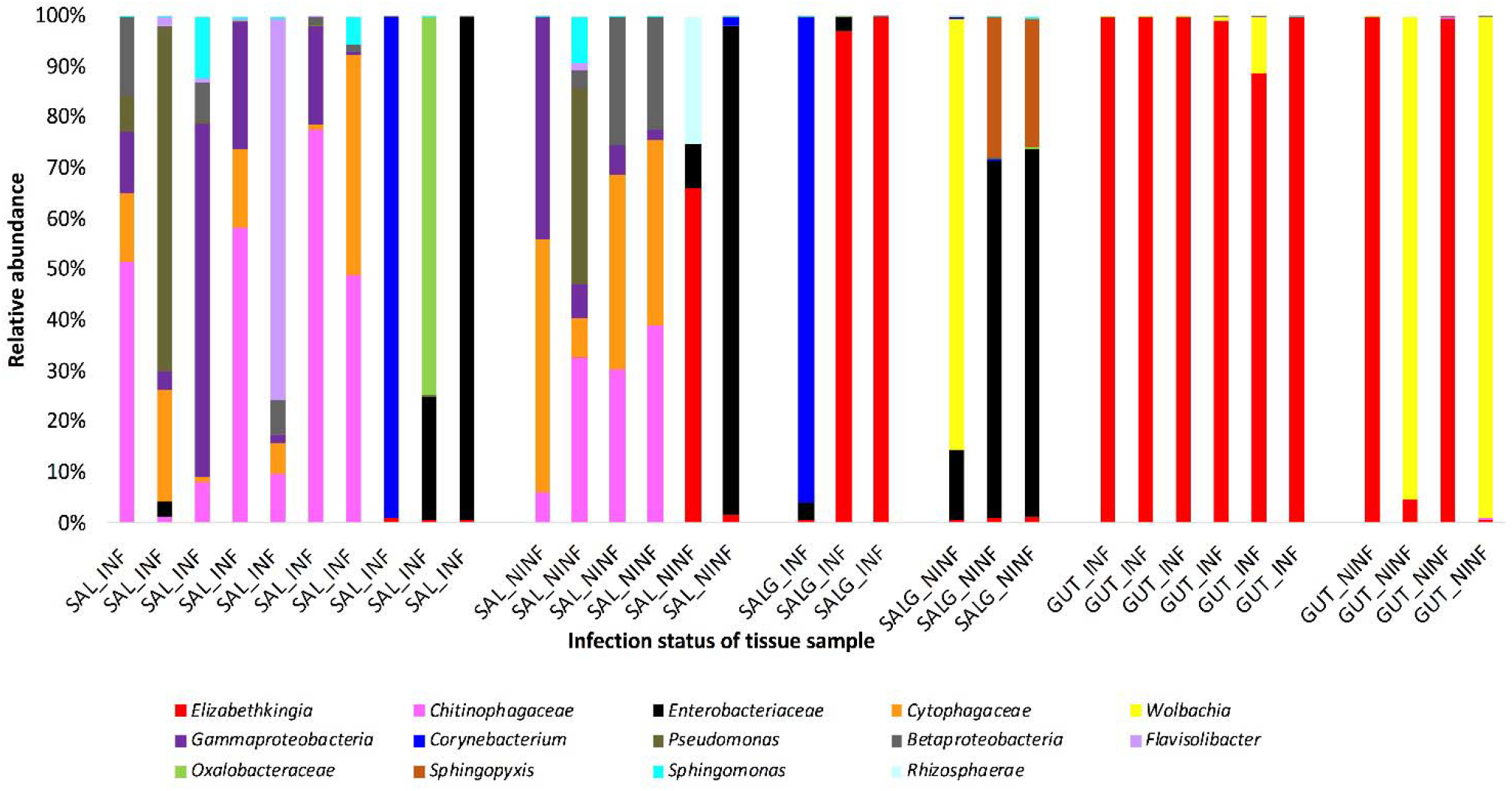
Relative abundance of bacterial taxa in individual *Aedes* saliva, salivary gland and midgut samples. Illumina MiSeq 16S rRNA sequencing was used to identify bacterial taxa and distributions were visualized using a cut-off of 7000 reads. Each stacked bar plot illustrates the taxonomic distribution of bacteria identified in individual *Aedes* saliva, salivary gland or midgut samples. Samples were taken at 7 days post-feeding. Blood fed individuals were further separated into NINF (fed non-infectious blood meal [BM]) and INF (fed Zika virus-positive BM and acquired an infection). SAL_INF represents saliva samples obtained from individuals fed on INF BM and successfully transmitting the virus; SAL_NINF represent saliva samples obtained from individuals exposed to NINF BM; SALG_INF represent individual salivary gland samples exposed to INF BM and infected; SALG_NINF represent salivary glands from individuals fed on NINF BM; GUT_INF represent midgut samples exposed to INF BM and infected, and GUT_NINF represent midgut samples from individuals fed on NINF BM.

The microbial profile was diverse across tissues (PERMANOVA, F-value=0.9998, R^2^= 0.3111; P <0.001; Figure 2). Midguts were dominated by Bacteroidetes with a higher relative abundance identified in ZIKV infected midguts relative to the unexposed midguts. Salivary glands, on the other hand, were dominated by Proteobacteria. Surprisingly, bacterial taxa from mosquito saliva was much more diverse with a range of Bacteroidetes, Proteobacteria and Cyanobacteria (Chao1 [ANOVA], P < 0.001, F-value= 20.58). This analysis clearly demonstrates the dominance of specific genera in individual tissues. For instance, ZIKV infected midguts and salivary glands were overwhelmingly dominated by *Elizabethkingia*. Uninfected midguts were dominated by either *Elizabethkingia* or *Wolbachia*, yet salivary glands from uninfected individuals were dominated by Enterobacteriaceae (Figure 1).

**Figure 2.**
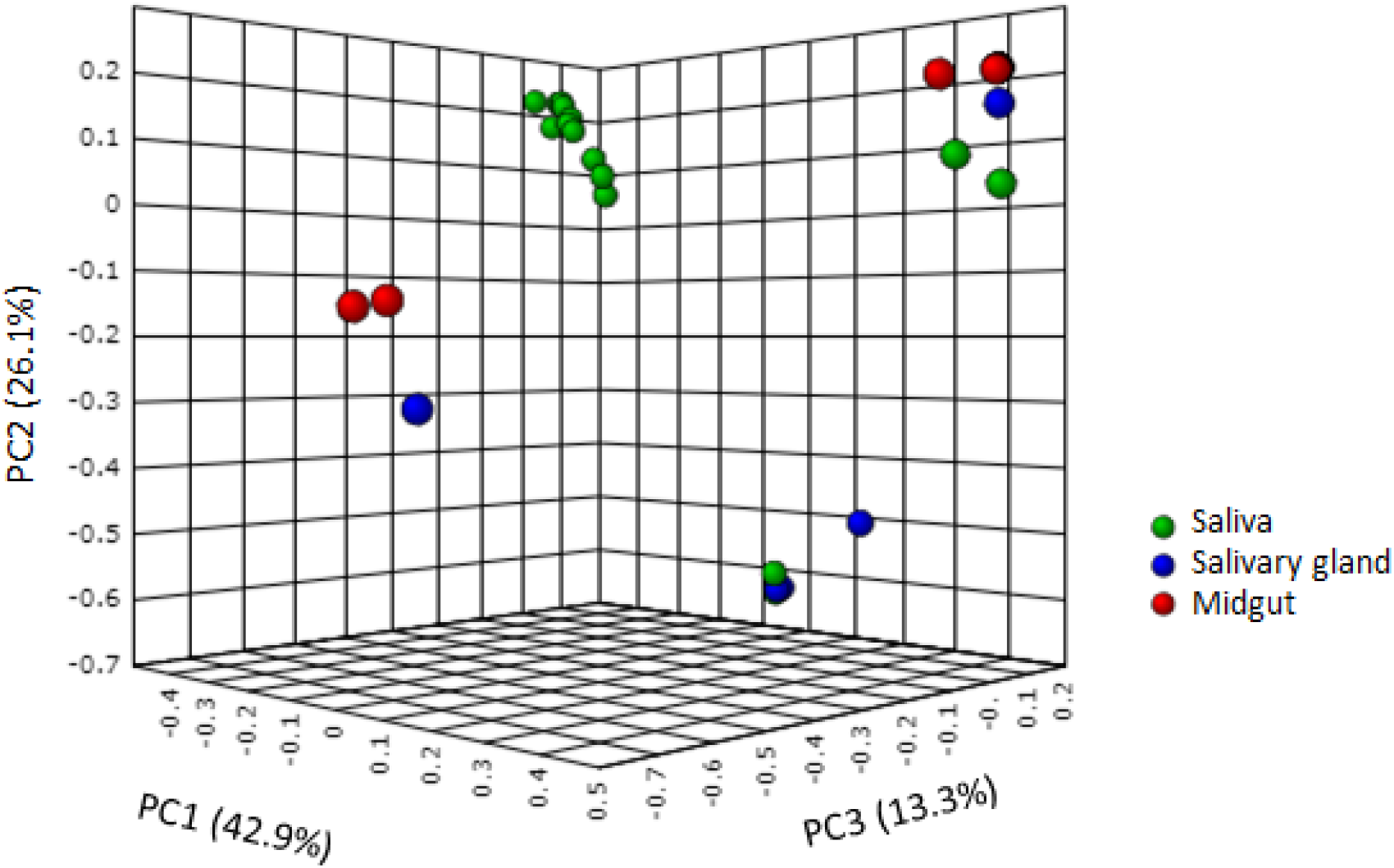
Tissue-specific diversity of microbes in *Ae. albopictus*. Principal component analyses were completed to assess the genetic relationships among and between bacterial taxa identified in different tissue types. Despite a number of shared individual bacterial genera, significant separation by tissue type was identified [PERMANOVA; F-value:6.9998; R-squared: 0.3111; P <0.001].

### *Aedes* saliva contains diverse microbiota including *Elizabethkingia anophelis*

The core microbes in the saliva included: *Pseudomonas, Cytophagaceae, Chitinophagaceae, Betaproteobacteria, Gammaproteobacteria, Comamonadaceae, Chryseobacterium, Flavisolibacter, Bradyrhizobiaceae, MLE1_12* and *Sphingomonas*.

The most abundant taxa present in the saliva (*Elizabethkingia, Chitinophagaceae, Enterobacteriaceae, Wolbachia, Pseudomonas* and *Betaproteobacteria*) were also present in the salivary glands and midgut samples (Figure 1).

We observed presence of taxa of public health importance in the saliva of *Ae. albopictus* exposed to infectious and non-infectious blood meals (Figure 1). These include *Elizabethkingia, Enterobacteriaceae* and *Pseudomonas*.

We employed a quality control measure to assess potential sources of saliva contamination. A cotton pad obtained from the container housing mosquitoes exposed to an infectious blood meal, a non-infectious blood meal, a sterile cotton pad, filter sterilized 10% sucrose solution used to feed mosquitoes, and filter sterilized mosquito diluent used to collect mosquito saliva, were incubated overnight in LB media. The control was sterile LB media. Bacterial growth was observed in the culture tube that had the cotton pad used to feed both mosquitoes exposed to the infectious blood meal as well as the non-infectious blood meal. No growth occurred in the culture tube with sterile cotton pad, sterile 10% sucrose solution, the mosquito diluent or the negative control. Introduction of bacteria during feeding by mosquitoes cannot be ruled out for taxa identified exclusively in saliva.

*Elizabethkingia* was present in both ZIKV -infected and uninfected midguts, salivary glands and saliva. On the other hand, *Enterobacteriaceae* and *Pseudomonas* were present in the saliva and salivary glands, but not midguts, of *Ae. albopictus* exposed to both ZIKV (infectious) and non-infectious blood meals (Figure 1; Figure 3).

**Figure 3.**
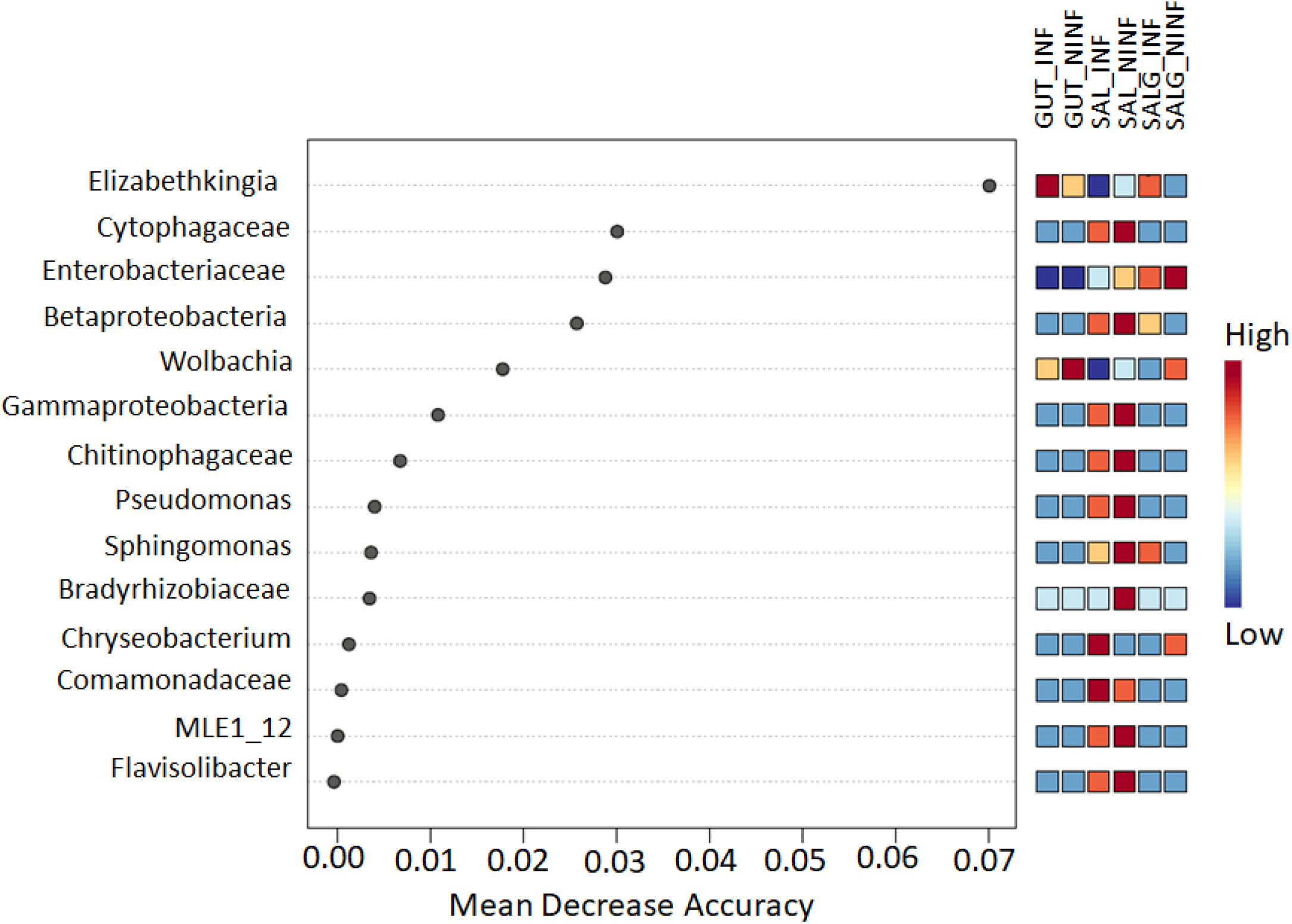
Association of bacterial taxa with both tissue and Zika virus infection status in *Ae. albopictus* mosquitoes. The Random forest classifier was used to determine the association of differentially abundant microbial genera with both tissue-type and Zika virus infection status of *Aedes* mosquitoes. Mean Decrease Accuracy is reported for each of the taxa. This measure is obtained by removing the relationship of a taxa and measuring increase in error. The taxa with highest mean decrease in accuracy is considered to have the highest association with the state. The highest mean decrease in accuracy was identified for *Elizabethkingia*. In addition, *Elizabethkingia* and *Enterobacteriaceae*, taxa associated with human disease, were found to be present in the midguts, saliva and salivary glands of *Aedes* mosquitoes exposed to both infectious (INF) and noninfectious (NINF) blood meals (BM).

### Relationship between *Elizabethkingia* and ZIKV in *Ae. albopictus*

The 16S *rRNA* amplicons of saliva, salivary glands and midgut samples were sequenced on the Illumina MiSeq platform. *Elizabethkingia* was identified in both saliva (10 out of 18) and salivary gland (6 out of 6) samples of *Ae. albopictus* exposed to non-infectious and infectious blood meals (Figure 1). *Elizabethkingia* sequences were also present in the midguts of *Ae. albopictus* exposed to infectious or non-infectious blood meal (10 out of 10) (Figure 1). Furthermore, a higher proportion of *Elizabethkingia* sequences were identified in saliva collected following non-infectious blood meals relative to the saliva collected following ZIKV infection (T-test; Mann Whitney test, P = 0.0256; Figure 3; Figure 4A). Conversely, while *Elizabethkingia* was identified in both the ZIKV-infected and uninfected *Aedes* salivary glands, ZIKV-infected salivary glands had a higher proportion of *Elizabethkingia* reads relative to their uninfected counterparts (F test, P< 0.0001; Figure 4A). Overall, the ZIKV - infected *Ae. albopictus* midguts had a higher relative amount of *Elizabethkingia* compared to other tissues (ANOVA; Kruskal-Wallis test, P = 0.0027; Figure 4A).

**Figure 4.**
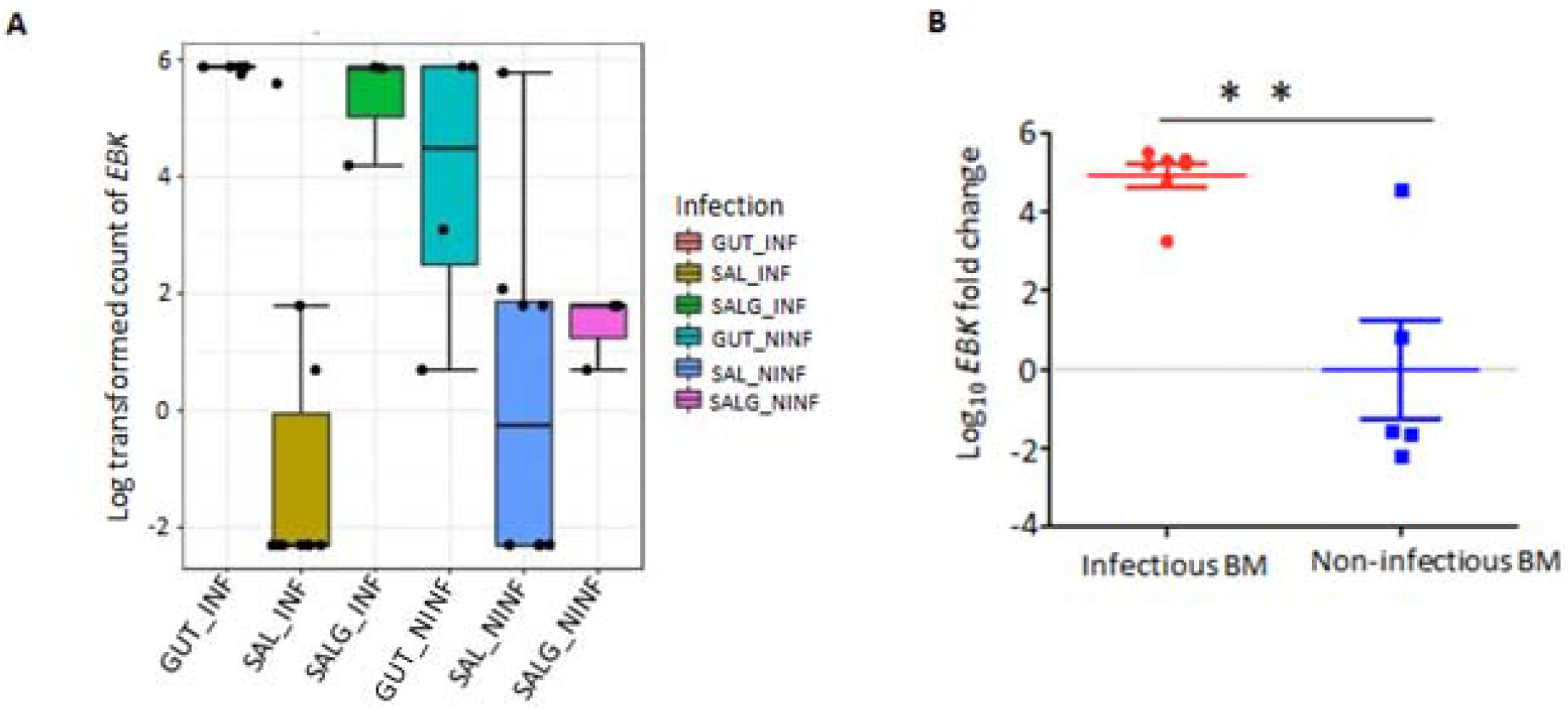
Relative *Elizabethkingia* levels in *Ae. albopictus*. **A. Tissue tropism.** Relative quantification of *Elizabethkingia* in the midgut, salivary gland and saliva of *Ae. albopictus*. Levels were tissue-specific and dependent on Zika virus (ZIKV) exposure. GUT_INF-infected gut; SAL_INF - infected saliva; SALG_INF - infected salivary gland; GUT_NINF – gut samples of *Ae. albopictus* exposed to NINF BM; SAL_NINF – saliva samples from *Ae. albopictus* exposed to NINF BM; SALG_NINF – salivary gland samples of *Ae. albopictus* exposed to NINF BM. **B. Relative levels of *Elizabethkingia* in Zika virus infected and uninfected *Ae. albopictus* midguts.** Infection of *Ae. albopictus* midguts by ZIKV (INF) was associated with higher *Elizabethkingia rpoB* transcript levels (T-test; Mann Whitney test P = 0.0051) relative to midguts following non-infectious blood meals (NINF BM) at 7 days post-feeding.

Although exposure of *Ae. albopictus* midguts to non-infectious blood meal resulted in an increase in the amount of *Elizabethkingia* in the midgut, Real-Time (RT) qPCR assay demonstrated a higher load of *Elizabethkingia* in the midguts exposed to ZIKV infectious blood meals as compared to the *Ae. albopictus* midguts exposed to non-infectious blood meals (T-test, Mann Whitney test, P = 0.0051; Figure 4B).

To further investigate the relationship between *Elizabethkingia* and ZIKV measured in *Ae. albopictus* midguts, *Elizabethkingia* bacterial loads were quantified together with ZIKV viral loads both *in vivo* (ZIKV-infected midguts, (N = 22) and *in vitro* (U4.4 cells co-infected with *Elizabethkingia* and ZIKV; three biological replicates, 1 biological replicate, n = 4 wells) using RT-qPCR.

Despite the fact that feeding on a ZIKV infectious blood meal was associated with higher levels of *Elizabethkingia* (Figure 4B), fold change in expression of the *Elizabethkingia rpoB* gene from ZIKV-infected *Ae. albopictus* (N = 22) was negatively correlated to ZIKV absolute quantities (Spearman r = −0.3985; P < 0.05; Figure 5A). ZIKV viral loads ranged from 3 - 11 log_10_ ZIKV RNA copies/ml, while fold change of *Elizabethkingia* ranged from −1.8 −2.2 log_10_ CFU/ml. To clarify the potential for direct interaction between *Elizabethkingia* and ZIKV, we performed an *in vitro* assay on the *Ae. albopictus* derived U4.4 cell line. Cells were co-infected with *Elizabethkingia* and ZIKV or infected with ZIKV only, and viral loads were compared. At 2 dpi, we measured high amounts of ZIKV in the control, ZIKV/*Elizabethkingia* OD 1.0 (8.5 Logs CFU/ ml) and ZIKV/ *Elizabethkingia* OD0.5 (8.0 Logs CFU/ml) co-infected cells while the ZIKV/ *Elizabethkingia* OD0.2 (7.7 Logs CFU/ml) co-infected cells had significantly lower ZIKV titers (ANOVA; Kruskal-Wallis test, P = 0.0070). On the other hand, at 4 dpi, we measured a significant reduction in ZIKV titers in all ZIKV/*Elizabethkingia* co-infected U4.4 cells relative to the control (ANOVA, Kruskal-Wallis test, P = 0.0079; Figure 5B).

**Figure 5.**
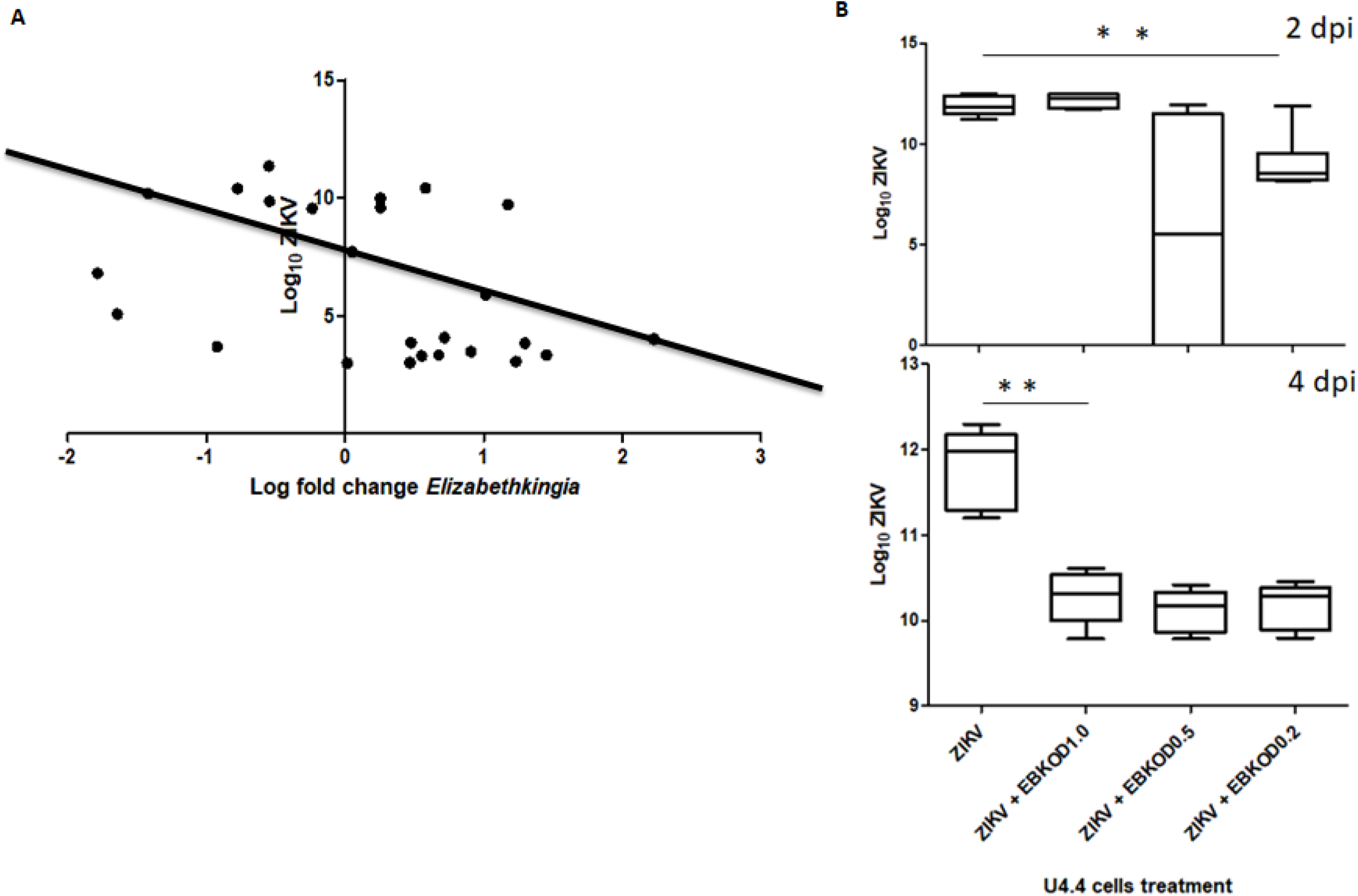
Relationship between *Elizabethkingia* and Zika virus proliferation. **A. *In vivo* comparison in *Ae. albopictus* midguts.** *Aedes albopictus* were infected with a bloodmeal containing 8.3 log_10_ PFU/ml ZIKV and of both expression of *Elizabethkingia rpoB* gene and ZIKV levels were assessed at 7 days post infection. The RT qPCR results were normalized using S7 gene. The X axis represents the log fold change of *Elizabethkingia*, while Y axis represents log_10_ ZIKV PFU/midgut. A negative correlation between ZIKV and *Elizabethkingia* was demonstrated (Spearman r = −0.3985; P < 0.05). **B. *In-vitro* assay of *Elizabethkingia* and Zika virus in U4.4 cells** U4.4 cells were infected with 8.3 log_10_ PFU/ml ZIKV following 2 days exposure to *Elizabethkingia* (EBK, experimental) or media alone (control). Experimental wells were previously exposed to either OD 1.0 (8.5 log_10_ CFU/ml EBK), OD 0.5 (8.0 log_10_ CFU/ml EBK) or OD 0.2 (7.7 log_10_ CFU/ml EBK). At both 2 and 4 days post infection (dpi) ZIKV levels were significantly higher in the ZIKV only controls relative to the EBK co-infected cells (ZIKV + *Elizabethkingia;* ANOVA; Kruskal-Wallis test, **P <0.01).

### Phylogenetic analysis of *Elizabethkingia*

To taxonomically classify the *Elizabethkingia* strain that we identified we sequenced the *Elizabethkingia rpoB* gene amplified from the cDNA of the *Ae. albopictus* salivary gland. The phylogenetic analysis demonstrated that the *Elizabethkingia anophelis* strain identified in our study has a 100% sequence identity to *Elizabethkingia anophelis* which was originally isolated from the midgut of *Anopheles gambiae* [20]. This strain has a 99% identity to *Elizabethkingia endophytica*, originally isolated from sweet corn [21]. It was closely related to the *Elizabethkingia genomosp*. 4 [22], *Elizabethkingia miricola*, isolated from condensation water in space station [23] and *Elizabethkingia bruuniana*[24]. *E. anophelis* has a 92% identity to *Elizabethkingia meningosepticum*, originally isolated from human cerebrospinal fluid [25] (Figure 6).

**Figure 6.**
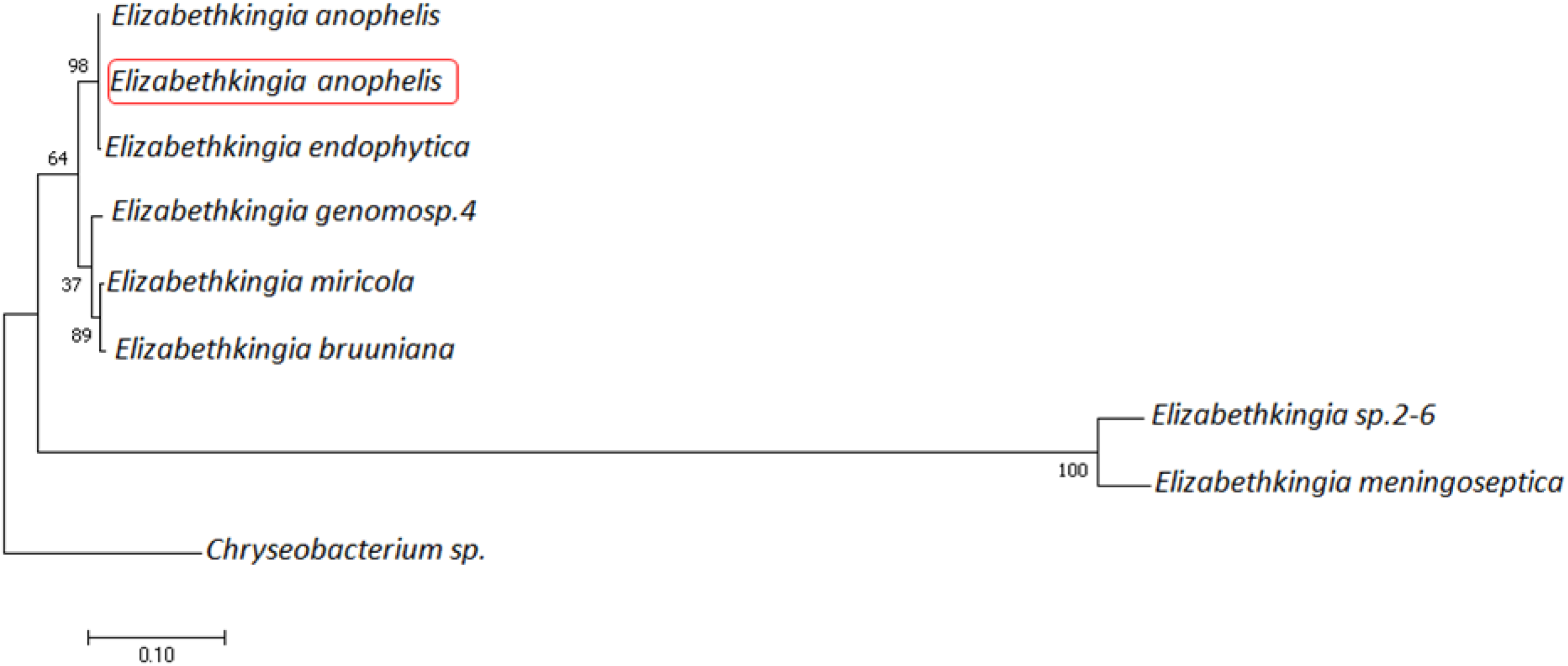
Phylogenetic analysis of *Elizabethkingia rpoB* gene. Phylogenetic affiliation of *Elizabethkingia* taxa from the salivary gland of *Ae. albopictus* based on a partial sequence of the *rpoB* gene (GenBank Accession number: MT096047). Phylogenetic dendrograms were constructed in MEGA. Bootstrap proportions are shown on branches. *Chryseobacterium* was used as an outgroup.

## Discussion

This study has demonstrated *Elizabethkingia* in the saliva of *Ae. albopictus* mosquitoes. Further, phylogenetic analysis has identified the strain of *Elizabethkingia* isolated from an *Ae. albopictus* salivary gland to have 100% sequence identity to *Elizabethkingia anophelis*, first isolated from the midgut of the mosquito *Anopheles gambiae*[26]. The *Elizabethkingia* genus was previously found to be the dominant bacterial genera in the salivary glands of both *An. gambiae* and *An. coluzii* [27]. To the best of our knowledge, this is the first study to demonstrate *E. anophelis* in the saliva of *Aedes* mosquitoes. Our findings bear clinical significance as *E. anophelis* has been identified as a cause of sepsis in adults and children, as well as a cause of neonatal meningitis [9]. By 2016, *E. anophelis* had caused an outbreak in Central African Republic [6], Singapore [7] and two outbreaks in the Midwest, United States that resulted in morbidity and mortality of ^~^ 30% [8–10]. The Midwest outbreak occurred primarily in community settings [8] and despite extensive investigations, no point source of infection has been identified in either the Midwest or Singapore outbreaks [9,10]. While there is currently no evidence of mosquito-borne transmission with any of these outbreaks, our data demonstrate the potential for involvement of mosquitoes in the maintenance of *E. anophelis* and other bacterial pathogens.

The identification of *Elizabethkingia* in the saliva of *Ae. albopictus* infected with ZIKV necessitates the investigation of the impact of *Elizabethkingia* on transmission efficiency of ZIKV and other arboviruses, and additionally on how co-transmission of bacteria could influence the early events, progression and severity of viral infections in the host. Our results present a new dimension into the vector-host interface dynamics. Several molecules present in mosquito saliva have been shown to suppress the host inflammatory or immune responses [28–33] thus enhancing dissemination [34]. Specific salivary proteins such as biogenic amine-binding D7 protein [35], saliva serine protease CLIPA3 [36], salivary factor LTRIN [37] and saliva protein AgBR1[38] modulate vertebrate infection. However, a gap in knowledge exists on the contributory role of mosquito salivary microbes in potentiating pathogenicity or transmission efficiency in vertebrate hosts. Salivary microbes may provide crucial targets for the control of disease-causing pathogen transmitted by mosquitoes.

In addition to *Elizabethkingia*, we observed a proliferation of other taxa associated with human disease throughout *Aedes* mosquitoes following exposure to a blood meal. This included bacteria belonging to the family Enterobacteriaceae, which have been shown to be opportunistic pathogens [39–41]. Enterobacteriaceae are gram negative bacteria that includes *Escherichia coli, Klebsiella, Salmonella, Shigella* and *Yersinia pestis. Enterobacter* spp., which we identified as among the primary bacteria associated with *Ae. albopictus*. These bacteria are known to produce molecules with hemolytic activity [42,43] and the increase in abundance of Enterobacteriaceae in the saliva of *Ae. albopictus* may therefore be an evolutionary mechanism that exploits blood digestion to ensure their horizontal transfer as well as infection of new hosts.

*Wolbachia* was identified in the saliva, salivary gland and midguts of *Ae. albopictus* in this study. *Wolbachia* is an intracellular endosymbiont primarily transmitted vertically through the female germline[44,45]. It can also undergo transfer between somatic and germ line cells. Horizontal transfer independent of cell to cell contact utilizing host cell phagocytic and clathrin/dynamin-dependent endocytic machinery is possible [46]. Our results provide further proof of the possibility of horizontal transfer of *Wolbachia* through saliva. *Wolbachia* has been identified in the phloem vessels and in some novel “reservoir” spherules along the phloem after *Wolbachia*-infected whiteflies fed on cotton leaves [47]. These findings could have important implications for the exploitation of *Wolbachia* as a biological control of tropical diseases.

A significant reduction in *Ae. albopictus* midgut microbial diversity resulting from ZIKV infection was evident, yet the effect on individual genera was highly variable. Available evidence demonstrates that commensal microbiota regulate and are in turn regulated by invading viruses [48–60]. Viral infection often causes disturbance of the commensal host microbiota, resulting in dysbiosis and subsequently affecting viral infectivity of the host [50,51,56,61–65]. The dysbiosis of the commensal microbiota due to virus infection is mainly due to modulation of the immune responses by the virus resulting in immune suppression [66,67]. Our results are consistent with the findings of a study on the response of the total gut microbiota diversity of *A. japonicus* and *A. triseriatus* to infection with La Crosse virus, which revealed higher midgut bacterial richness among the midguts obtained from newly emerged individuals and lower richness among midguts exposed to viral infection or a non-infectious blood meals [50]. In addition, our study corroborates previous findings with West Nile virus infection on the bacterial diversity of *Culex pipiens*, which demonstrated that lower diversity was associated with virus exposure[51].

It is also notable that *Elizabethkingia* and *Wolbachia* appeared to have an exclusive presence in the midgut of the *Ae. albopictus* mosquitoes. It is possible that there is competitive exclusion of *Wolbachia* in the midgut by *Elizabethkingia*. Studies have shown that *Ae. albopictus* and *Cx. quinquefasciatus* naturally infected by both *Asaia* and *Wolbachia* have a co-exclusion relationship[51]. In species uninfected by *Wolbachia* such as *An. gambiae, An. stephensi* and *Ae. aegypti, Asaia* is capable of colonizing reproductive organs and salivary glands [68]. The predominance of *E. meningosepticum* in the *An. stephensi* midgut was shown to be due to its antimicrobial properties suppressing the growth of other midgut bacteria [69].

We demonstrated a significant negative correlation between ZIKV and *Elizabethkingia* as well as a reduction in the ZIKV titer in U4.4 cells co-infected with *Elizabethkingia* and ZIKV. Similarly, it has been demonstrated that *E. meningoseptica* metabolites obtained from the midgut of *Anopheles stephensi* exhibited antiparasitic effect on *P. falciparum* parasites *in vitro* and demonstrated *in vitro* toxicity on gametocytes[69]. We also measured a significant increase in *Elizabethkingia* bacterial load in the midgut infected with ZIKV relative to those exposed to a non-infectious blood meal. These findings corroborate the results of a previous study aimed at understanding the effect of ZIKV on the dynamic bacterial community harbored by *Ae. aegypti* [70], which demonstrated that the Flavobacteriacae family was the most abundant bacterial taxa. This was particularly the case among the ZIKV-exposed blood fed individuals, yet Flavobacteriacae decreased in abundance among ZIKV-exposed gravid individuals [70]. The interactions between bacteria and viruses are clearly complex. The mechanisms of interaction mostly benefit virus infiltration into cells directly by virus binding to a bacterial cell or viral utilization of a bacterial product [71]. Indirect mechanisms include: virus-induced increase of bacterial receptor concentrations; virus damage to underlying epithelial cells; virus displacement of commensal bacteria and virus suppression of the host immune system[71]. In the direct interactions, the commensal bacteria are considered capable of outcompeting viral pathogens and limiting tissue accessibility by competing for receptor binding sites reducing the binding of pathogenic bacteria or viral attachment. This may potentially explain the negative correlation that we observed in *Ae. albopictus* midguts. Our *In vitro* results which demonstrate decreased viral loads with *Elizabethkingia* co-infection, particularly at day 4 post-infection, corroborate previous studies [70] and clearly demonstrate the capacity for this bacterium to independently suppress viral replication and/or persistence. Further studies delineating the mechanism that *Elizabethkingia* utilizes to inhibit viral fitness in *Ae. albopictus* are needed, yet these data suggest the identification of a novel transmission barrier that could ultimately be exploited in the design of novel control measures.

Overall, to the best of our knowledge, this is the first study to demonstrate and characterize the microbiome of the saliva of mosquitoes. These data may have significant implications on the pathogen - host interface and our understanding of the impact of these microbes on the severity of disease in the vertebrate host. We have also demonstrated that the salivary gland and saliva of *Aedes* mosquitoes exposed to infectious or non-infectious bloodmeals contain the bacterial pathogen *Elizabethkingia anophelis*. Finally, our study has demonstrated both by *in vivo* and *in vitro* studies that *Elizabethkingia* and ZIKV are negatively correlated in the mosquito midgut. These findings together have implications for our understanding of the interaction of bacterial and viral agents in mosquitoes that could ultimately improve our understanding of disease epidemiology and inform novel control measures.

## Materials and methods

### Virus

This study utilized frozen virus sample. The ZIKV strain was HND (2016-19563, GenBank accession no. KX906952).

### Insect sampling and DNA preparation

*Ae. albopictus* were collected from Suffolk County, NYS in 2015 (kindly provided by I. Rochlin) and colonized at the New York State Department of Health (NYSDOH) Arbovirus Laboratory. F15 *Ae. albopictus* were vacuum-hatched and maintained at 27°C under standard rearing conditions. At eight days post mating, female adults were orally exposed to blood meals. The blood meal consisted of either 1:1 dilution of defibrinated sheep blood plus 2.5% sucrose; sodium bicarbonate (to adjust pH to 8.0) and virus, or a non-infectious blood meal containing a final concentration of 2.5% sucrose solution. Infectious blood meals contained 8.3 log_10_ PFU/ml ZIKV HND [72]. The female mosquitoes were exposed to blood meals in a 37°C pre-heated Hemotek membrane feeding system (Discovery Workshops, Acrington, UK) with a porcine sausage casing membrane. After an hour, the mosquitoes were anaesthetized with CO_2_ and immobilized on a pre-chilled tray connected to 100% CO_2_. Engorged females were separated and placed in three separate 0.6 L cardboard cartons (30 individuals per carton). In addition, 1 ml of each blood meal was transferred to a 1.5 ml Safe Seal microtube (Eppendorf, Hamburg Germany) and stored at −80°C. Blood fed females were maintained on 10% sucrose solution provided *ad libitum*.

At 14 dpi, the female mosquitoes were immobilized using triethylamine (Sigma Aldrich, St. Louis, MO, USA). To examine for ZIKV dissemination in the mosquito, the legs were removed from the mosquitoes before dissecting the gut and salivary gland from every individual. Saliva was collected under sterile conditions by inserting the proboscis of the female mosquitoes into a capillary tube containing ^~^20 μl fetal bovine serum plus 50% sucrose 1:1 for 30 minutes and subsequently ejecting the mixture into 125 μl Mosquito Diluent (MD; 20% heat-inactivated fetal bovine serum in Dulbecco phosphate-buffered saline plus 50 μg/ml penicillin/streptomycin, 50 μg/ml gentamicin and 2 μg/ml Fungizone (Sigma Aldrich, St. Louis, MO, USA). The midgut was dissected from individual mosquitoes and rinsed twice in sterile PBS before transfer to sterile microcentrifuge tubes and storage at −80°C until tested. Mosquito carcass and legs were then placed in individual tubes containing 500 μl MD and a bead. All samples were held at −80°C until assayed. A ZIKV-specific quantitative PCR assay that targets the NS1 region was utilized to obtain viral titer [73] from the legs, carcass, saliva, salivary glands and midguts as described by [72].

As a precautionary measure to prevent contamination of saliva, all the reagents used during the rearing and blood feeding of mosquitoes were filter sterilized and cotton pads were autoclaved.

To test for potential contamination during rearing, we incubated overnight (in LB media) a piece of cotton pad obtained from the gallon housing mosquitoes exposed to an infectious blood meal, mosquitoes exposed to a non-infectious blood meal, a sterile cotton pad not exposed to mosquitoes, filter sterilized 10% sucrose solution that was used to feed the mosquitoes, filter sterilized mosquito diluent, and sterile LB Media used as a control.

### Sequencing and analysis of microbiome of *Aedes* mosquito saliva, salivary glands and midguts

To determine the microbial profile in the *Ae. albopictus* saliva, salivary glands and midguts of each individual mosquito, midgut, legs, carcass and saliva samples were screened for virus. The individuals that were successfully infected and or transmitting the virus were identified. The cDNA was prepared from the RNA samples using iScript cDNA kit (Biorad, Hercules, California). To amplify starting material for downstream analysis, the saliva, salivary gland and midgut samples were individually whole transcriptome amplified according to the REPLI-g WTA single cell kit protocol (Hilden, Germany). The resulting amplified cDNA was diluted 1:100. Thereafter, 16S *rRNA* amplicon were amplified using the 16S primer set and conditions in Table 1.

**Table 1.**
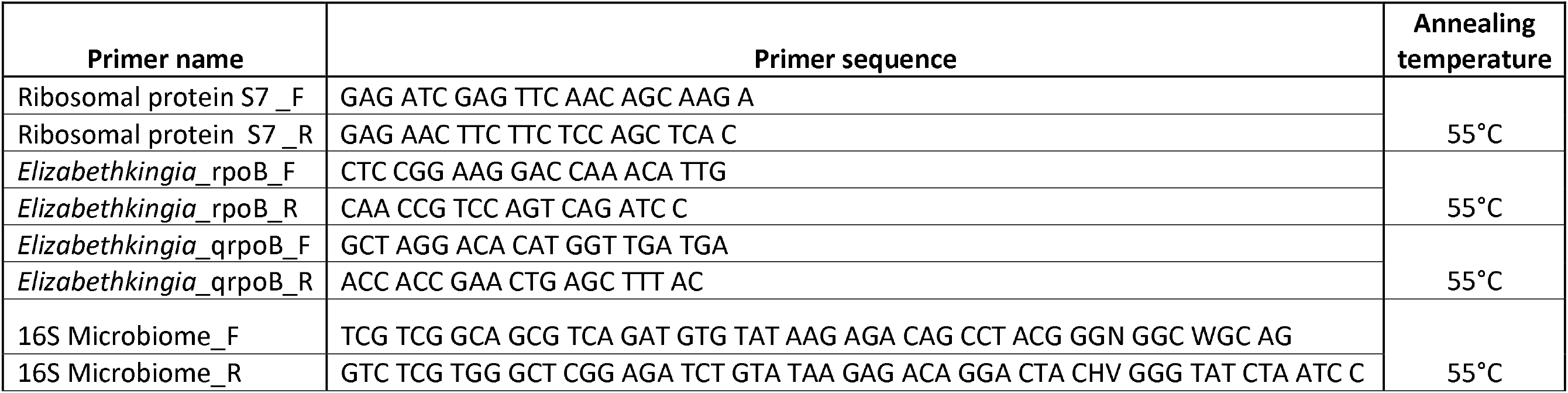
List of Primers used in this study.

The barcoded high-throughput amplicon sequencing of the bacterial 16S V3-V4 hypervariable *rRNA* was done at Wadsworth Center Applied Genomics Core (WCAGC). PCR reactions were carried out in a total volume of 50 μl, 5μM of 16S *rRNA* V3-V4 hypervariable region primers consisting of an Illumina barcode (Table 1) [51], 2 μl WTA cDNA, 8 μl deionized filter sterilized water and 36 μl AccuStart II PCR supermix (Quanta biosciences, Beverly, USA). To ensure that any potential contamination either at the PCR stage or at the sequencing step would be identified, two negative controls: water and non-template control were included at the PCR step and sequenced as well as a positive spike included during the sequencing run. A fragment size of ^~^460 bp of each sample was submitted to the WCAGC for sequencing together with the negative controls. Automated cluster generation and paired-end sequencing (250-bp reads) was performed on the Illumina MiSeq 500 cycle.

A total of 37 samples [6 infected midguts (INF MG); 4 non-infected midguts (NINF MG); 12 infected saliva (INF SAL); 6 non-infected saliva (NINF SAL); 3 infected salivary glands (INF SALG); 3 non-infected salivary gland (NINF SALG); 1 negative control (NC); 1 non-template control (NTC) and 1 positive spike (PC) were included in this study.

Analysis of the data was carried out on QIITA (https://qiita.ucsd.edu/) and MicrobiomeAnalyst [74]. In summary, to convert the de-multiplexed FASTQ files to the format used by QIITA, the command split libraries FASTQ was used. The clustering of the sequences into operational taxonomic units (OTUs), was done by the utilization of a closed-reference OTU picking based on the GreenGenes 16S reference database which matches the sequences based on a 97% sequence identity threshold before assigning a taxonomy. A BIOM-formatted OTU table was subsequently generated, to reduce the potential of alpha and beta diversity biases, the data was rarefied to a depth of 30 000 reads per sample. Finally, to visualize the taxonomic profiles of each sample, taxa bar plots were generated using the rarefied data artifact. Following the generation of the relative quantity of each taxon, we generated a taxa bar plot based on taxa with sequences ≥ 4 000.

The reads have been deposited in NCBI GenBank Short Read Archives with accession numbers: PRJNA608507, PRJNA608505, PRJNA608503, PRJNA608501, PRJNA608500, PRJNA608499, PRJNA608472, PRJNA608496, PRJNA608495, PRJNA608493, PRJNA608492, PRJNA608491, PRJNA608489, PRJNA608483, PRJNA608471, PRJNA608482, PRJNA608470, PRJNA608456, PRJNA608450, PRJNA608481, PRJNA608449, PRJNA608448, PRJNA608480, PRJNA608479, PRJNA608477, PRJNA608437, PRJNA608434, PRJNA608427, PRJNA608428, PRJNA608429, PRJNA608375, PRJNA608426, PRJNA608424, PRJNA608416, PRJNA608418, PRJNA608422, PRJNA608436 and PRJNA608419.

The alpha diversity profiling (the number of species of taxa in a single sample) was calculated using the Chao1 diversity measure and significance testing was done using the ANOVA method in Microbiome Analyst. Chao1 index estimates the richness of taxa in a sample by extrapolating out the number of rare organisms that may have been omitted due to under sampling. The index assumes that the number of observations for a taxon has a Poisson distribution and hence corrects for variance [75].

Beta diversity, a comparison of microbial community composition between two samples or two conditions, was analyzed by utilizing the Bray-Curtis dissimilarity statistic [76]. The distance matrix generated after comparing each sample to the other was visualized for dissimilarity between samples using the Principle Coordinate Analysis ordination method.

### Clustering and Biomarker analysis

Patterns of relative abundance of taxa species in response to different experimental factors was analyzed in Microbiome Analyst [74]. The distance measure was Euclidean [77] and the clustering algorithm utilized was Ward, an agglomerative hierarchical clustering procedure [78]. The Euclidean distance, a measure of the true straight-line distance between two points in Euclidean space, examines the root of square differences between coordinates of a pair of objects.

To provide an estimate of the most important taxa for classification of data resulting from different experimental factors, biomarker analysis was carried out using the random forest algorithm within Microbiome Analyst [74]. The default setting of the number of trees to grow and number of predictors to try was applied (500 and 7, respectively) with the randomness setting left on. Random forest algorithm [79] is a supervised classification algorithm of trees created by using bootstrap samples while training data and random feature selection in tree induction. It is an ensemble of unpruned classification or regression trees trained with the bagging method [80].

### Quantification of *Elizabethkingia* and Zika virus

Both *in vitro* and *in vivo* analyses to quantify the *Elizabethkingia* loads were performed using the primer pairs *rpoB* gene specific to *Elizabethkingia* using the PCR conditions described in Table 1. The ribosomal protein *S7* gene was utilized to normalize transcript levels (Table 1). Absolute quantity of ZIKV in individual midguts was obtained by isolating RNA using Trizol (ThermoFisher scientific, Massachusetts, USA). A ZIKV-specific quantitative PCR assay that targets the NS1 region was used for detection [73] and serially diluted ZIKV of known quantities was used to generate the standard curve and extrapolate viral load.

### Phylogenetic analysis of the *Elizabethkingia* isolated from *Ae. albopictus* salivary gland

In order to validate the identity of *Elizabethkingia* taxa identified in the *Ae. albopictus* salivary gland and saliva samples, *rpoB* gene (GenBank Accession number: MT096047) was amplified from the salivary gland cDNA using the *rpoB* primer pairs and PCR conditions as described in Table 1. The 558 bp PCR amplicon was excised from Agarose gel and cleaned using QIAquick gel extraction kit (Qiagen, UK). The amplicons were Sanger sequenced at the WCAGC.

The sequences were manually trimmed using Bioedit v7.1.9 and aligned using MUSCLE [81,82]. Upon trimming the primer region and the low-quality regions of the sequence, the FASTA sequence of the respective genes were blasted against the NCBI database. A total of 9 gene sequences published with the following GenBank accession numbers: CP023010.2, KY587659.1, CP016377.1, KY587657.1, CP035811.1, CP035809.1, CP011059.1, CP039929.1 and MN327643.1 were added to the phylogenetic analysis.

To construct the *Elizabethkingia rpoB* phylogenetic tree, Molecular Evolutionary Genetic Analysis (MEGA) software [83] was utilized. The phylogenetic tree was constructed using the Maximum Likelihood statistical method [84] with nucleotide substitution type.

The test of phylogeny was by bootstrap method [85] with 1000 bootstrap replications. The nucleotide substitution model was the Tamura 3-parameter model [86]. The rates among sites were uniform rates while the Heuristic method was the Nearest-Neighbor-Interchange.

### *In vitro* assay of *Elizabethkingia* and ZIKV co-infection

In order to study the growth kinetics of ZIKV co-infected with *Elizabethkingia*, U4.4 cells (*Ae. albopictus*), were co-infected with *Elizabethkingia* (kindly provided by Steven Blaire, The Connecticut Agricultural Experiment Station) and ZIKV.

*Elizabethkingia* bacterial strain was streaked on Luria-Bertani (LB) plates and incubated in a 10% CO_2_ incubator at 28°C overnight. A single colony was picked using a sterile plastic loop and used to inoculate 500 ml of sterile LB broth. The inoculated broth media was incubated overnight at 28°C in an orbital shaker shaking at 225 rpm. The *Elizabethkingia* overnight culture was subsequently diluted to OD 1.0 (8.5 log_10_ CFU/ml), OD 0.5 (8.0 log_10_ CFU/ml) and OD 0.2 (7.7 log_10_ CFU/ml).

Individual wells were seeded with 3 ml of 2 X 10^5^ U4.4 cells/ml suspended in M&M medium without FBS and incubated at 28°C for 1 h before adding FBS to each well at a final concentration of 20%. The cells were then incubated for 5 days in a 10% CO_2_ incubator at 28°C. After 5 days, supernatant was removed carefully, and the plates were infected with 100 μl of *Elizabethkingia* at OD 1.0 (8.5 log_10_ CFU/ml), OD 0.5(8.0 log_10_ CFU/ml) and OD 0.2 (7.7 log_10_ CFU/ml) for three biological replicates. The negative control plates were infected with 100 μl of 8.3 log_10_ PFU/ml. After 1 h of incubation at 28°C, 3 ml of M&M media containing 20% fetal bovine serum (FBS) were added to each well and incubated at 28°C.

To determine whether the U4.4 cells were successfully infected with *Elizabethkingia*, a single well from each replicate of the different dilutions of *Elizabethkingia* infection was harvested at 2 dpi.

The cells were spun at 1000 rpm and washed three times with sterile phosphate buffered saline (PBS). After the third wash, 100 μl of the washed cells were used to inoculate 3 ml of LB media and incubated overnight at 28°C in an orbital shaker at 225 rpm.

The experimental wells were infected with 100 μl of 8.3 log_10_ PFU/ml ZIKV after a 2 day incubation with *Elizabethkingia*. Cells which were co-infected with *Elizabethkingia* and ZIKV, as well as the ZIKV only control, were harvested at both 2 dpi and 4 dpi. RNA was extracted from the cells using Trizol (ThermoFisher scientific, Massachusetts, USA). The ZIKV quantities were measured as previously described.

### Statistical analysis

Statistical analysis was performed with GraphPad Prism version 5.0. Differences in *Elizabethkingia* quantities were assessed using two-tailed T test. Viral loads were compared using an ANOVA test. Finally, the *Elizabethkingia* /ZIKV correlation analysis was carried out using Pearson tests.

## Acknowledgements

This publication was supported by the Cooperative Agreement Number U01CK000509 funded by the Center for Disease Control and Prevention. Its content is solely the responsibility of the authors and do not necessarily represent the official views of the Center for Disease Control and Prevention or the Department of Health and Human Services. These studies were also partially funded by the National Institutes of Health award R21AI131683. We would like to extend our gratitude to Illia Rochlin of Suffolk County Health Department for providing us with *Ae. albopictus* mosquitoes and the Arbovirus laboratory insectary crew for mosquito maintenance. We additionally thank the Wadsworth Center Applied Genomics core for sequencing and the Wadsworth Center Tissue and Media facility for providing cells and media.

## Author Contributions

MGO: Involved in the study design, conducted the experiments, analyzed the results and drafted the paper

PA: Trained me in the cell culture, was involved in the antibiotic treatment of the Aedes mosquitoes and assisted in mosquito manipulation exercises

SJ: Involved in the mosquito rearing, assisted in mosquito manipulation exercises

DC: Assisted in mosquito manipulation exercises

KL: Generated and propagated the ZIKV infectious clone utilized in this study

KLD: Involved in the study design, participated in the writing of the paper

CAT: Involved in the study design, participated in the writing of the paper

## Materials and correspondence

Correspondence and material requests should be addressed to CAT

